# The impact of global transcriptional regulation on bacterial gene order

**DOI:** 10.1101/405894

**Authors:** Pablo Yubero, Juan F. Poyatos

## Abstract

Bacterial gene expression depends on the allocation of limited transcriptional resources provided a particular growth rate and growth condition. Early studies in a few genes suggested this global regulation to generate a unifying hyperbolic expression pattern. Here, we developed a large-scale method that generalizes these experiments to quantify the response to growth of over 700 genes that *a priori* do not exhibit any specific control. We distinguish a core subset following a promoter-specific hyperbolic response. Within this group, we sort genes with regard to their responsiveness to the global regulatory program to show that those with a particularly sensitive linear response are located near the origin of replication. We then find evidence that this genomic architecture is biologically significant by examining position conservation of *E. coli* genes in 100 bacteria. The response to the transcriptional resources of the cell results consequently in an additional feature contributing to bacterial genome organization.

## Introduction

Transcription regulation is one of the fundamental mechanisms by which bacteria adapt gene expression to changing environmental conditions. Apart from the specific action mediated by transcription factors (TFs), expression is modulated by a global regulatory program determined by the physiological condition of the cell. Initial studies correlated this condition to the availability of core constituents of the expression machinery: free RNA polymerase, tRNAs, ribosomes, etc. (1, 2), but many other interacting components can play a role such as cell volume, or the alarmone (p)ppGpp (3–5). These works also provided an effective protocol to describe the influence of all these elements: the dependence on physiology was linked to growth rate at exponential phase independent of the particular nutrients fixing that rate. Therefore, the examination of the global program reduced to the quantification of expression response to changes in growth rate and growing conditions.

That global physiology complements specific regulation matters in many aspects. Indeed, growth rate dependencies can interfere directly with genetic circuits and change their operation, for example, by shifting the bistability regime of a switch or allowing for different antibiotic resistance strategies (6, 7). Costs of synthetic genetic circuits on cell physiology and the consequences of the latter on the function of the circuits made the subject also relevant in applied areas, e.g., Synthetic Biology (8). Beyond “simple” genetic circuits, the interplay of global regulation and cell resource allocation can modify many essential features at the system level (9, 10). In fact, mechanistic approaches revealed that the global regulatory program contributes to determining fundamental trade-offs involving the finiteness of the cellular size, energy, and ribosomal fraction (11).

To examine the activity of this program, the choice of constitutive genes as the primary model is clear: promoters of these genes lack any interaction with specific DNA-binding TFs, and thus, they are *a priori* constantly available for transcription initiation. Therefore, constitutive genes are subject only to physiological regulation. An alternative approach is to mutate the TF binding sites of non-constitutive genes to assess the separate (mutant) and combined (wild type) effect of global and specific regulation. Studies applying these approximations included however only a few genes (12–14), and thus we lack a large-scale evaluation of global transcriptional regulation.

Beyond its evaluation, it is also intriguing to examine to what degree global regulation could impact bacterial genomic organization, as it is the case for specific regulation (15). One of the factors contributing to this regulation is copy number as gene dosage depends on the growth rate and on the distance to the origin of replication *oriC* of the chromosome. This is due to the overlap of multiple replication rounds at fast growth rates (multifork replication). Indeed, the position and copy number of ribosomal genes in *Escherichia coli* are tuned to maintain fast growth rates (16). We could nevertheless ask if the global transcriptional regulation excluding the copy number affects genomic architecture. At least two scenarios can be postulated. One in which promoters that are intrinsically sensitive to the global transcriptional regulation, i.e., excluding copy number contribution, are located far from *oriC* to compensate for the small, almost negligible, increase in copy number at large growth rates. And a second scenario, where those promoters are located near *oriC* to further enhance their activity with growth rate. In the first situation, the influence of the global transcriptional program is compensated along the chromosome whereas in the second, copy number strengthens the dependence between expression and growth rate. Either solution would reveal design principles of genome architecture.

In this work, we introduce a procedure that enables us to first examine at a large scale the response of ~700 genes (with no known explicit regulation by TFs) to the separate and combined effect of the global regulation program and the chromosomal copy number variations due to multifork replication in *E. coli*. To this end, we develop a method that uses experimental time-series of growth rate and promoter activity of a fluorescent reporter library (17, 18), which has proven to be one of the best tools to study *in vivo* gene expression at large scale. This allowed us to recognize a core set of strictly constitutive genes presenting a promoter-dependent hyperbolic response. For these genes, we quantify the most sensitive to the global program and observe that they are significantly located near the origin of replication. This presents the proximity to *oriC* as an important feature to enhance the control of expression and suggests that the location of these genes could be particularly conserved in species in which this control is desirable, e.g., those experiencing faster or more variable growth rates. We examine evidence in this respect with the analysis of the correlation between position conservation of the corresponding *E. coli* genes in 100 bacterial species and the number of replication rounds, maximal growth rate, and environmental variability of the species’ habitat.

## Results

### Quantifying chromosomal promoter activity at a large scale

To quantify the promoter activity of chromosomal genes (PA_*chr*_) we developed a method that makes use of promoter activity measurements obtained with low-copy plasmids (PA_*pl*_). This is of particular interest as the availability of a fluorescent library in *E. coli* (17) could then be used to determine PA_*chr*_ at a large scale while reducing the experimental burden of locating expression reporters on the chromosome. We build upon a previous gene expression model in which the promoter activity measured is proportional to the promoter activity per gene copy (pa) and the gene copy number per cell (*g*) and that is inversely proportional to cell volume (v) (6).

We first decouple the copy number signal of the plasmid *g_pl_* that contributes to PA_*pl*_ (Fig 1A). Since the replication of these plasmids is synced to the end of the cell cycle (19, 20), the plasmid copy number *g_pl_* is proportional to that of terminal regions in the chromosome (*g_ter_*). Moreover, in the context of this fluorescent library, earlier experimental results showed that proportionality between *g_pl_* and *g_ter_* is equal to 5 independently of both growth-rate (up to 1.8~dbl/h) and measurement approach (balanced growth and time-series) (13). We can thus consider Cooper and Helmstetter’s model (21) describing the copy number of a chromosomal gene *g_chr_* for a given growth rate μ (*g_chr_* = 2^*μ*[*C*(1−*m*)+*D*]^, *m* represents the normalized distance to the origin of replication of the gene) to obtain the plasmid copy number: *g_ter_* ∝ *g_pl_* = 5 · 2^*μD*^, the values of *C* and *D* are obtained by interpolation from experimental measurements (22).

Second, we decouple the growth-rate dependent cell size v(μ). The cell size law reads v=2^*μ*(*C+D*)^ in units of unit cell size, and it robustly predicts cell volume under several perturbations (23). Multiple studies have shown that cell volume is proportional to cell mass, which is in turn well captured by optical density measurements (24, 25).

**Fig. 1.**
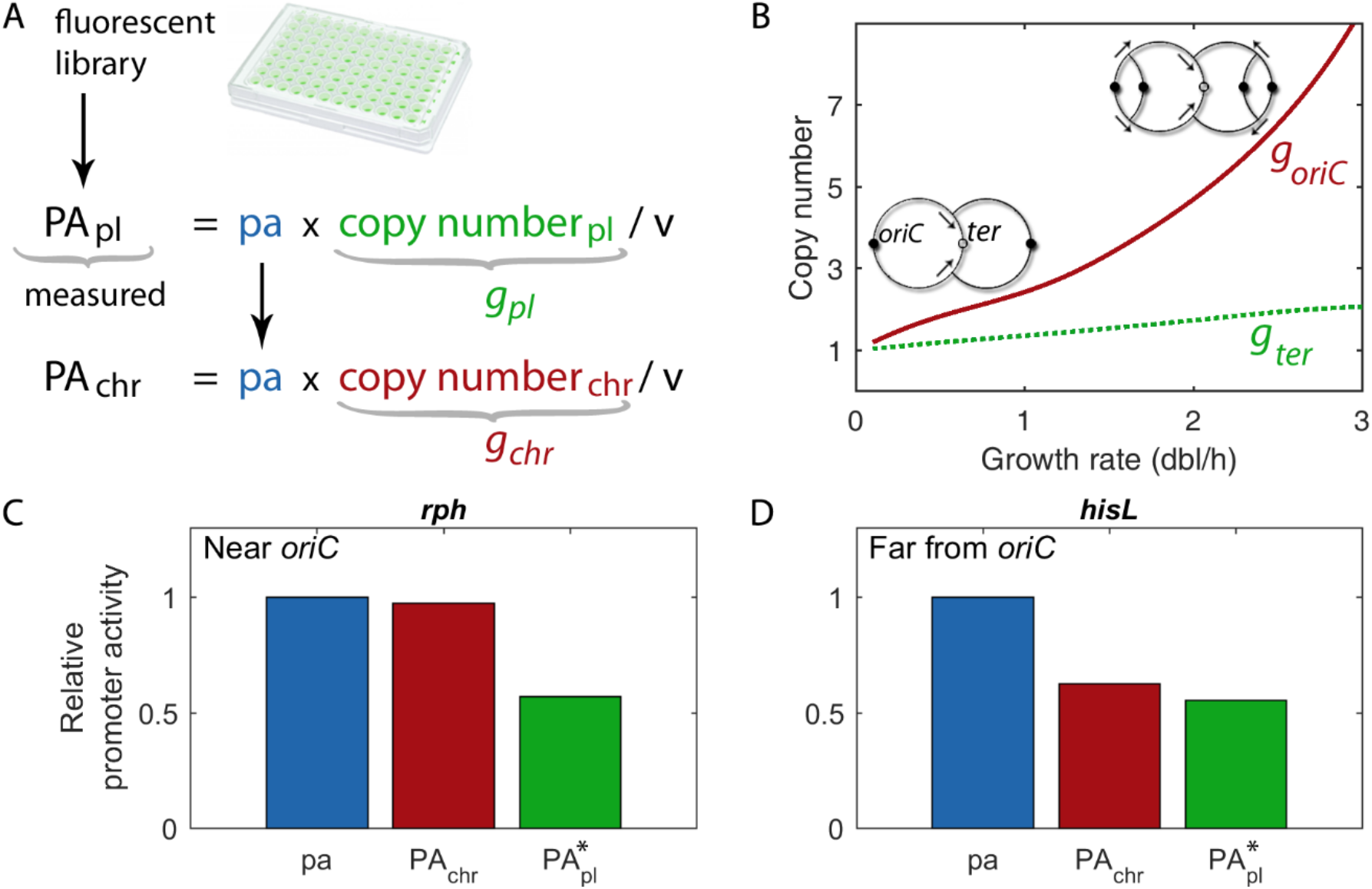
Decoupling promoter activity from gene copy number. (**A**) Promoter activity per single gene copy, pa, can be obtained from experimental data of promoter activity quantified with a plasmid library, PA_*pl*_, once the plasmid copy number *g_pl_* and the growth-rate dependence of the cell volume, v, are known. With this, one can calculate the promoter activity of a chromosomal gene, PA_*chr*_, by using Cooper and Helmstetter’s model (Materials and methods). (**B**) Chromosomal multifork replication makes the copy number per cell of chromosomal genes *g_chr_* dependent on both growth rate and gene location in the chromosome. At a faster growth rate, the number of origins of replication *oriCs* (red solid line and black dots in sketch) increases due to the overlap in time of multiple replication rounds. Arrows show the direction of replication forks. In the case of plasmids with low copy number, as the one used in the plasmid library (pSC101), *g_pl_* is proportional to the number of terminal regions (ters) in the cell (green dotted line). (**C,D**) Relative differences in promoter activity (pa, PA_*chr*_, PA_*pl*_) for two genes at different chromosomal locations for a fixed growth rate (normalized to the corresponding pa). Genes (*rph* and *hisL*) are located at distances *m_rph_* =0.04 and *m_hisL_* =0.80 from *oriC*. Observe that chromosomal promoter activity depends strongly on the location of the gene. Data was obtained in balanced growth at a rate μ~0.9 dbl/h (18). For comparability, we show PA*_*pl*_=PA_*pl*_/5 to normalize for the proportionality constant between *g_ter_* and *g_pl_* (see Main text for details).

This enabled us to compute promoter activity per gene copy, pa = PA_*pl*_ v / *g_pl_*, where the effect of gene copy number and volume is excluded, and chromosomal promoter activity PA_*chr*_ = PA_*pl*_ *g_chr_* / *g_pl_* where both effects are included (Fig. 1A-B).

Figures 1C-D show the resulting promoter activities of two example cases using experimental data from (18); genes *rph* and *hisL* located at distances from the origin of replication of *m_rph_* =0.04 and *m_hisL_*=0.80, respectively. Differences in chromosomal promoter activity become relevant when comparing genes at different positions in the chromosome. In this way, the distinction between the promoter activity per gene copy (pa, Fig. 1A) and chromosomal promoter activity (PA_*chr*_, Fig. 1A) emphasizes the added effects of multifork replication depending on the location of the gene. Note that, due to the increase in copy number, the level of PA_*chr*_ of promoters closest to oriC keeps up with the increase in cell volume.

### Constitutive genes show a promoter-specific hyperbolic response to global regulation

We applied the previous approach to characterize the global program at a large scale. Constitutive genes appear as the most suitable model given the absence of any specific regulation acting on them, and a list of these genes can be proposed with the information available in current databases (Materials and methods and also Discussion). However, characterizing the response of constitutive promoters in a traditional manner, i.e., from balanced growth measurements in different carbon sources, limits the scalability of the approach. We follow then here an alternative method and consider instead measurements of promoter activity during dynamic changes of growth rate in a specific carbon source. Note that these measures, in the case of constitutively expressed genes, correlate well with those observed under balanced growth in different growth media (13).

We thus processed the time series data of the set of 708 “constitutive” genes of *E. coli* included in the fluorescent library (18) (Materials and methods). Instead of measuring hundreds of genes in many distinct carbon sources, we considered data during exponential and late-exponential growth (within the first 5hrs) in glucose medium supplemented with amino acids to obtain profiles of instantaneous promoter activity and growth rate; PA_*pl*_(*μ*) profiles (Fig. S1 in supplemental material, Materials and methods). Data derived in this way can be decoupled from their plasmid context in order to get chromosomal, PA_*chr*_ (*μ*), and per gene, pa(*μ*), profiles (previous section).

After computing PA_*chr*_ (*μ*) we applied a clustering algorithm that grouped all resulting profiles into four classes (Materials and methods; cophenetic correlation coefficient c=0.80, Fig. 2A,B). Class 1 corresponds to promoters whose activity increases following the expected Michaelis-Menten profile with distinct parameters (Data set S1 in supplemental material), as it is expected from earlier works (4, 13, 14), whereas classes 2 and 3 correspond to promoter activities that decreases or remain mostly constant across growth rates, respectively. Finally, class 4 includes promoters with a non-monotonic profile that has maximum promoter activity at intermediate growth rates. These classes are robust whether PA_*chr*_ (μ) or pa(μ) profiles are used for the classification (Fig. S2 and Data set S1 in supplemental material).

**Fig. 2.**
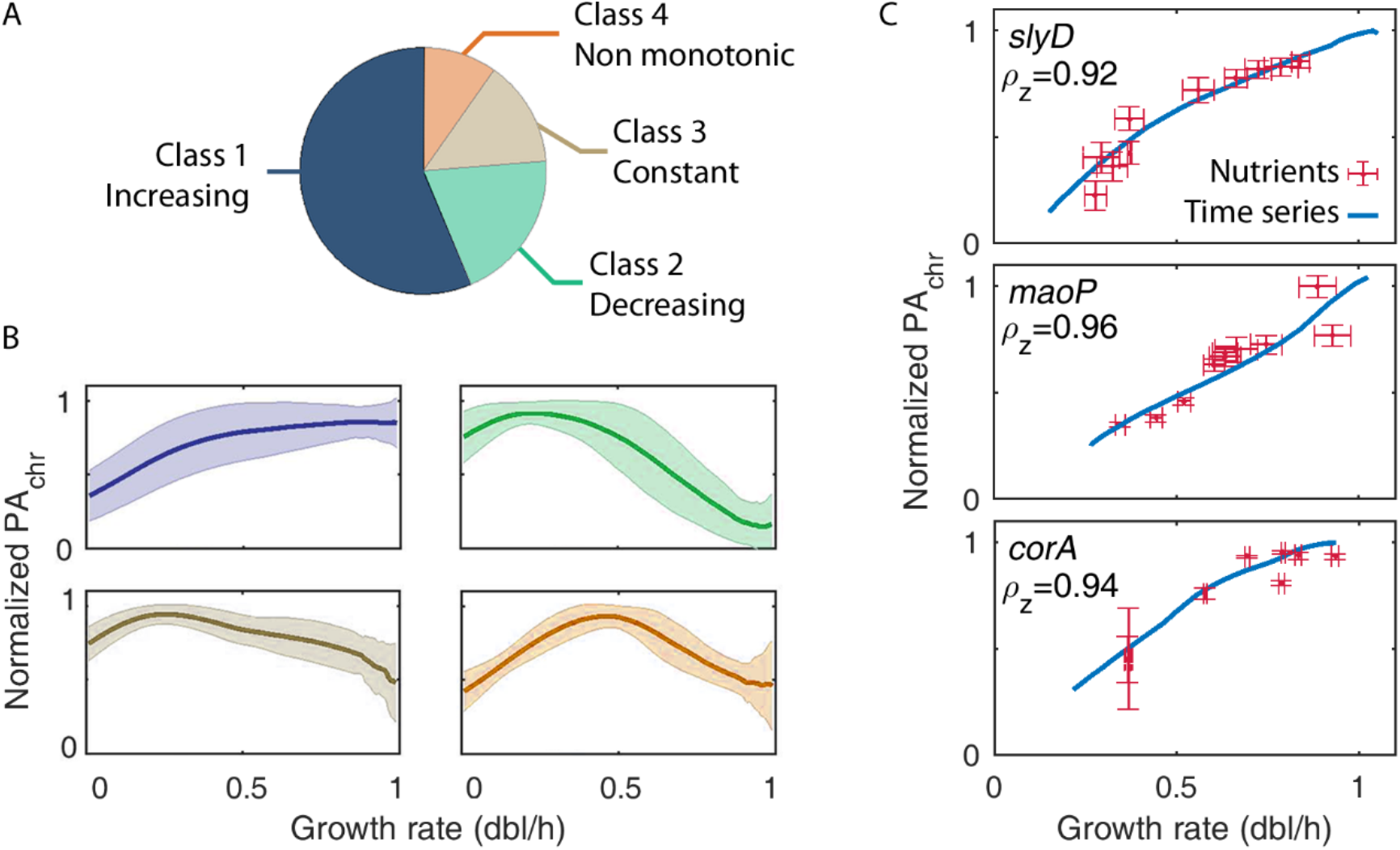
The clustering algorithm groups the PA_*chr*_(*μ*) profiles into four classes, of which only the first could be validated experimentally. (**A**) Fraction of promoters found in each of the four classes. Using a clustering algorithm, we grouped the PA_*chr*_(μ) profiles of about 700 genes with no known TF regulation into four classes following their growth-rate dependency. Only class 1 comprises profiles with the expected behavior from earlier works. (**B**) Mean profile of each class (solid line) and one standard deviation (shaded). Note that these profiles were obtained from time series on a single growth medium (Materials and methods). (**C**) Experimental measurements of PAchr from balanced growth in ten different media (red crosses, mean and sd from three replicates) validate our approach of inferring the profiles from time series data in glucose supplemented with amino acids (blue solid line, Materials and Methods) of genes in class 1. We also find large linear correlations ρ_z_ between our own time series data and that of Zaslaver *et al*. (2009). Fig. S3 in supplemental material shows the experimental results for all 12 promoters tested (three from each class). Data of *corA* grown in glycerol and arabinose resulted in fluorescence levels below the background and are not shown.

To test the approach of inferring PA_*chr*_ profiles from time series on a single growth medium, we experimentally measured the promoter activity profiles PA_*chr*_ (μ) of 12 promoters –chosen among all four classes– from balanced growth data in 10 different growth media (Materials and methods, Fig. S3 in supplemental material). The method appeared only particularly robust for class 1 promoters (56%). Indeed, Figure 2C shows the experimental results of three genes within the first class *slyD, maoP* and *corA* (a brief description of these genes is available in Materials and methods). For genes in classes 2-4, not only do we not recover experimentally the cluster profiles, but we also fail to recover the expected Michaelis-Menten hyperbolic pattern of constitutive genes.

In addition, to verify if this lack of signal could be related to the reliability of the clustering algorithm, we added random noise to the chromosomal promoter activity profiles and measured the mean number of recovered genes to the original classification expressed in percentage (10 realizations, Materials and methods). For normally distributed relative levels of noise up to 10%, we recovered 93% of promoters assigned to class 1, whereas the rest of the classes had recovering rates between 35% and 79%. This suggest overall that the discrepancies that we find with classes 2 to 4 are not related to the approach itself, but rather that these promoters might experience some unknown specific regulatory mechanisms.

However, the robustness with which class 1 promoters are identified and characterized suggests that promoters in this class are likely constitutive. For this reason, we discard promoters from classes 2-4 in the following and we use only high confidence profiles from class 1. Moreover, these results also suggest that as observed previously, during the first hours after balanced growth and before stationary phase, their expression can be well determined by the physiological state of the cell (13).

### Promoters sensitive to global regulation are located closer to the origin of replication

Beyond the previous classification, we noted different genes within class 1 promoters with distinct sensitivity to the global regulation. To quantify sensitivity, we fitted PA_*chr*_(μ) profiles to a Michaelis-Menten equation:

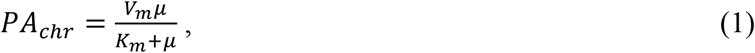

where *V_m_* is the maximum promoter activity and *K_m_* is the growth rate at which PA_*chr*_ (*μ*) is half maximal; note that μ records the global program, and that the different responses emphasize a promoter-specific rather than an unspecific pattern (4, 9, 13). Next, we classified as linear (Fig. 3A) or saturable (Fig. 3B) profiles those with *K_m_* > 3 dbl/h and 0.1<*K_m_* < 3 dbl/h, respectively (Materials and methods). The classification is robust with respect to different thresholds within realistic growth rates, only a small number of genes, around 7% for both pa and PA_*chr*_, have *K_m_* values within 2 and 4 dbl/h.

**Fig. 3.**
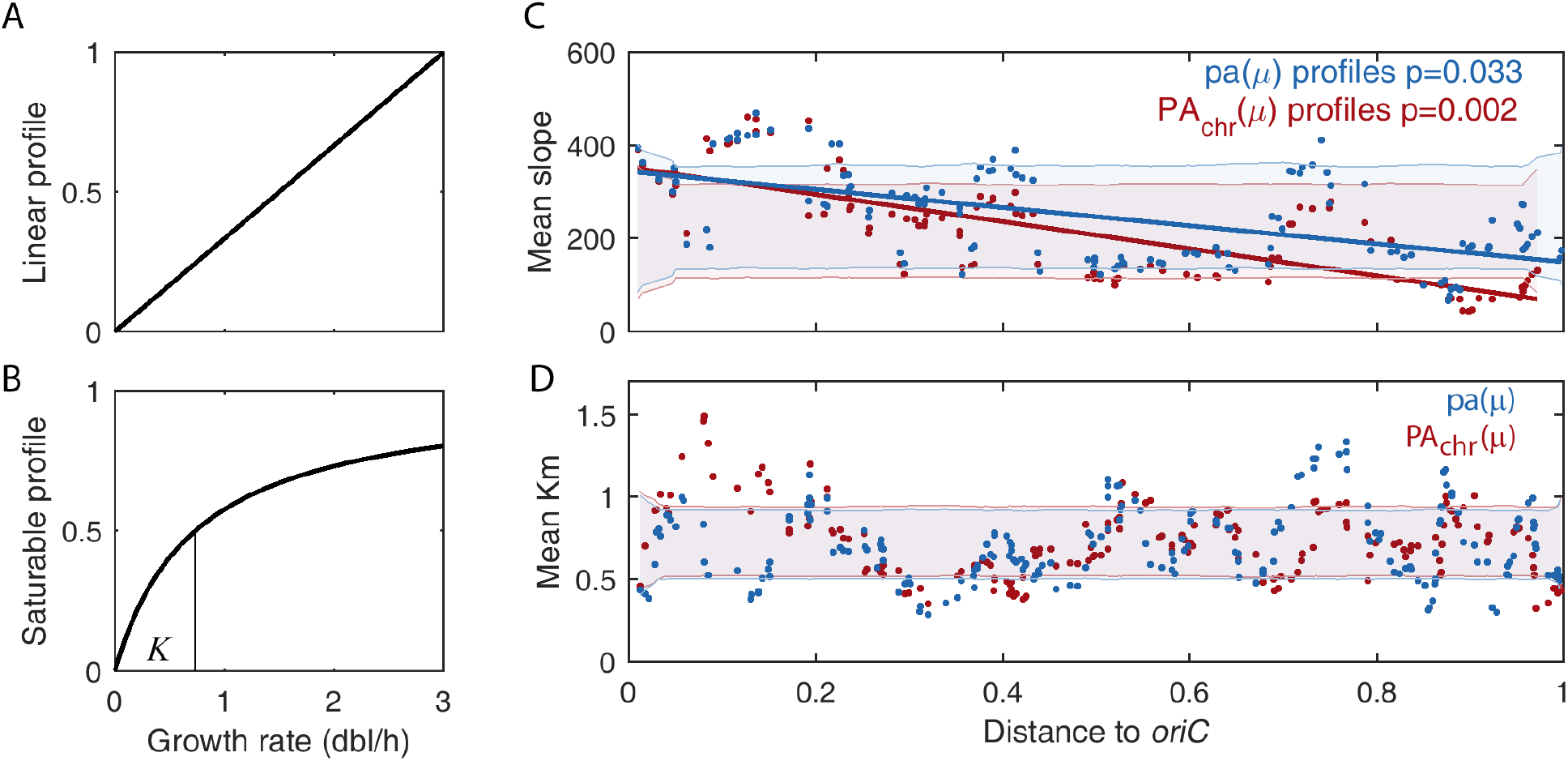
Promoters that are most sensitive to growth rate are located closer to the origin of replication. Two profiles of promoter activity can be identified: linear (**A**) and saturable (**B**). Sensitivity to the global program is proxied by the slope in the case of linear profiles, and the growth rate at which activity is half maximal (*K_m_*) in the case of saturable profiles. (**C, D**) Running averages of the sensitivity to the global program of promoters with linear and saturable profiles respectively of PA_*chr*_ (red dots) and pa (blue dots). The sensitivity of linear profiles decreases linearly with the distance to *oriC* (solid lines); this pattern is only significantly observed in saturable profiles when considering PA_*chr*_. Shading denotes one standard deviation of sensitivities obtained from a permutation test with 10^4^ randomizations.

In the case of promoters with linear profiles, we defined the sensitivity to the global program as the slope of the PA_*chr*_ (μ) profile, such that larger values stand for larger increases in promoter activity for fixed changes in growth rate. In the case of saturable promoters, we took *K_m_* as a proxy of their sensitivity to the global program: for smaller values of *K_m_* the promoter activity becomes near saturation at smaller growth rates, thus becoming less sensitive to changes in growth rate. We also computed sensitivities of pa(μ) profiles determined in an analogous manner (Materials and methods, Fig. S4 and Data set S1 in supplemental material).

We then asked if there exists an association between sensitivity and chromosomal location, given that one of the factors that influence these responses is multifork replication, relevant near *oriC*. Figure 3C-D shows the running average of the sensitivities to the global program along the chromosome of constitutive promoters with linear and saturable profiles, when including and excluding the effects of multifork replication, i.e., PA_*chr*_ (*μ*) and pa(μ), respectively. We observed that the sensitivity of linear profiles decreases linearly with the distance to *oriC* more abruptly and more significantly when considering PA_*chr*_ than pa. In the case of saturable constitutive promoters, we notice that only when considering PA_*chr*_ there is a significant peak within *m*<0.20 of the chromosome (p<0.05).

In general, these results suggest that saturable promoters in *E. coli* are located across the genome independently of their promoter activity per gene copy. On the contrary, linear promoters that are most growth-rate dependent are preferentially located near the origin of replication where they can further boost their expression due to increased copy numbers at large growth rates.

### Global regulation acts as a gene position conservation force

In light of the previous results, it is reasonable to hypothesize that both modes of regulation (gene location, and the sensitivity to the global program) would act synergistically in those species experiencing multiple overlapping replication rounds, hence preserving gene order. Inversely, gene order should be lost only in species living in rather stable environments or experiencing long doubling times.

To evaluate this hypothesis, we examined next if these genes maintain their proximity to *oriC* in other species as a function of some characteristics of the species: their maximum growth rate, the variability of the environment where they live and, the capacity for multifork replication (a function of genome size and minimal doubling time). We performed a homolog search across 100 species to compute the corresponding chromosomal displacement (Fig. 4A, Materials and methods). Displacements of the half most growth-rate-dependent genes near *oriC* (*m*<0.2, Fig. 3) are compared against the null hypothesis, i.e., displacement is independent of sensitivity. This is scored by the probability of finding a larger mean displacement of gene groups of the same size chosen randomly among all constitutive promoters at *m*<0.2, for linear or saturated growth-rate dependencies (Fig. 4B). Smaller values of this score, termed the position conservation, represent non-conserved locations of promoters.

**Fig. 4.**
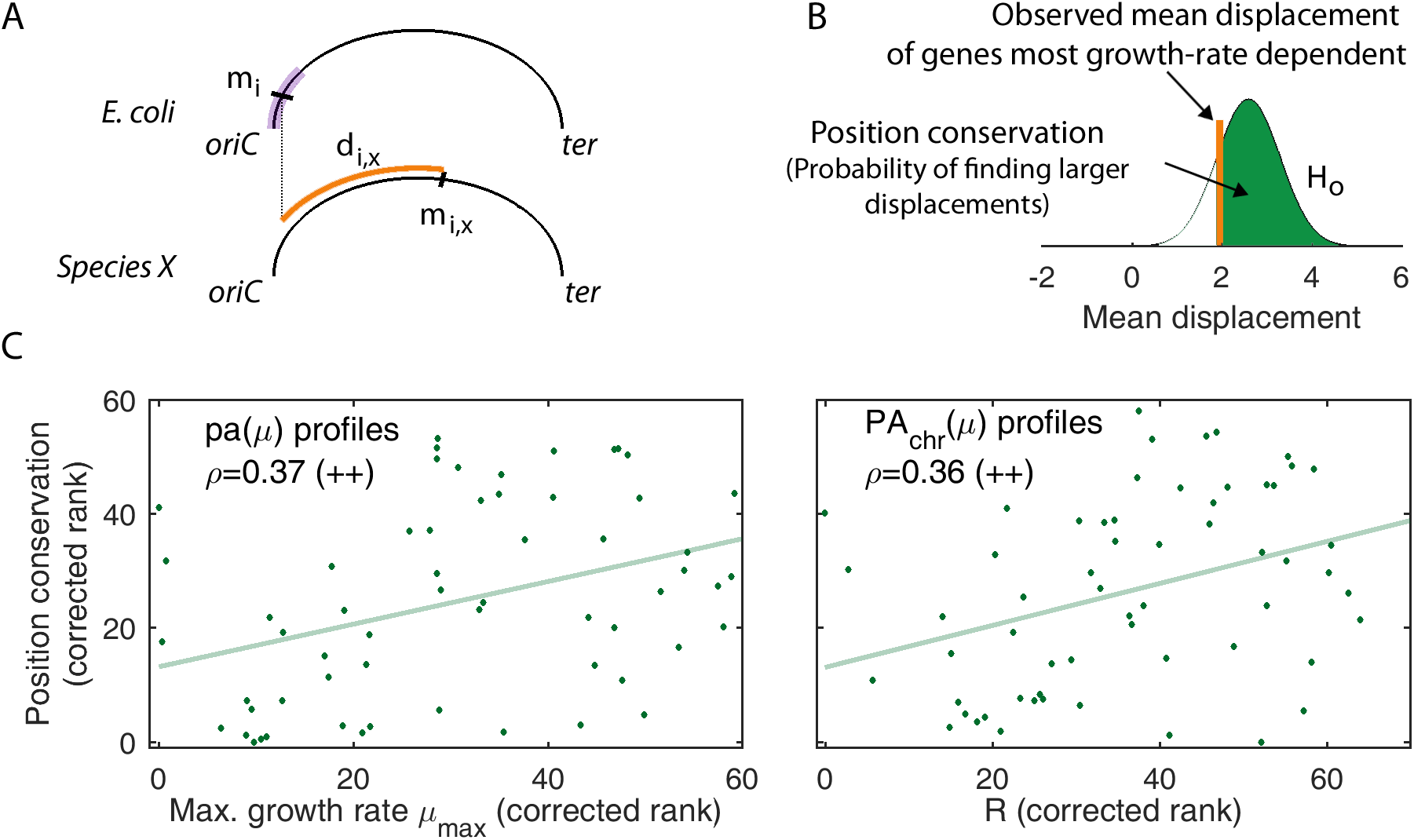
The position conservation of constitutive genes near *oriC* that are most dependent on the global program correlates with the maximum growth rate and R. (**A**) Gene’s position conservation is computed from the displacement of a gene in *E. coli* (*m_i_*) within the *m*<0.2 region (purple), with respect to its homolog in other species (*m_i,X_*). (**B**) In every species, the observed mean displacement of genes that are most dependent on the global program and are located at *m*=<0.20 is tested against the displacement of the rest of constitutive genes at *m*=<0.20 (H_o_). (**C**) The most predictive partial correlations (Spearman ρ, and light green solid line, both p<0.01) of the position conservation of the half most growth-rate-dependent lineal profiles near *oriC* in *E. coli* were obtained with R, for PA_*chr*_(μ) profiles, and the maximum growth rate, for pa(μ) profiles. Variables are corrected for phylogenetic inertia (Materials and methods).

We studied next the association between the position conservation and three main species features: environmental variability (env), relevance of multifork replication (R), and maximal growth rate (as the inverse of the minimal doubling time, μ_max_ = τ_min_^−1^). Environmental variability was based on an earlier environmental classification (26), while minimal doubling time with genome size was estimated to compute R; the ratio between the maximal chromosome’s replication time and the minimal doubling time as a measure of the importance of multifork replication effects for an organism (27). For each class of promoter dependence (linear and saturated, pa and PA_*chr*_) we measured the partial Spearman’s rank correlation ρ between the corresponding position conservation and env, R, or μ_max_ while controlling in all cases for phylogenetic distance (Materials and methods).

In the case of lineal promoters, we obtained a significant correlation between position conservation and R or μ_max_, respectively (all Spearman ρ with p<0.01). The numerical values of ρ are in line with those obtained in other gene order studies (27). Correlations with R are only slightly stronger when the global program includes the multifork effect (PA_*chr*_; 0.36 *vs*. 0.35), as expected from the definition of R, whereas maximal growth rate and R are equally relevant when not including the multifork dosage effect (pa; 0.37 *vs*. 0.37). Figure 4C explicitly shows these correlations; pa(μ) *vs*. maximum growth rate and PA_*chr*_(μ) *vs*. R, respectively. The position conservation of saturable promoters was not significant in any case.

Overall these results support our hypothesis that the impact of the global transcriptional program on gene order is a general feature of bacterial species, especially in those that undergo multifork replication. In fact, the position of genes exhibiting a particularly sensitive linear response tend to be conserved in species with larger values of R and fast growth rates.

## Discussion

We quantified growth-rate dependencies of over 700 prospective constitutive genes in *E. coli*, arguably the best gene collection to explore the effects of the physiological state of the cell, the global program, on gene expression. This is based on an approach that obtains promoter activity as if reporters had been inserted in the chromosome and that characterizes growth-rate dependencies from dynamical data. Both features reduce the costs and difficulties of large-scale experiments.

Half of the promoters that we examine present a Michaelis-Menten rate law confirming earlier reports (4, 13). That we verify this class with experiments in which the dependency is obtained using conventional approaches (growth rate being modified with the utilization of different carbon sources; data obtained at steady state) supports our approach. However, we also find three other patterns that differ. This does not seem associated with the method itself as our experimental characterization of these responses did not recover hyperbolic profiles. These genes could be perhaps subject to additional layers of regulation or other hidden structural aspects. However, we observed no signal of a particular enrichment on specific sigma factors or AT content in the promoter region or upstream of it (Fig. S5A-B in supplemental material), as large AT content is known to favor DNA bending and thus protein-DNA interactions (28, 29), in particular, upstream of the promoter region in the UP element (30). Additionally, although the supercoiling state of the chromosome is known to affect gene expression, no quantitative or even qualitative genome-wide regulatory model is yet available (31). We considered instead data on independent supercoiling macrodomains (32) to notice again no signal (Fig. S5C in supplemental material).

Overall, the success in predicting the response of over 50% of promoters with the original list (arguably the truly constitutive ones) demonstrates the significance of the global program beyond balanced growth (12). Within these promoters we distinguish subsets that reveal especially sensitive to growth rate and that are selectively located in the chromosome. Indeed, genes with either linear or saturable profiles show larger sensitivities to growth rate within 20% of the replichore closest to *oriC*. This pattern is partially maintained when we control for the multiple replication fork effect, i.e., when we consider pa(μ) profiles instead of PA_*chr*_(μ). We thus propose a model in which multifork effects and the global program (excluding gene copy) work in combination: promoters that are most growth-rate dependent in *E. coli* benefit from a larger increase in gene expression at large growth rates (Fig. S6 in supplemental material).

However, the fact that *E. coli* coordinates different mechanisms to obtain a multiplicative effect of enhanced expression of genes near *oriC* might not necessarily be a general property of bacterial genomes. This precise coupling might have been selected in bacteria for which multifork gene dosage fluctuations are relevant: those that are subject to variable growth rates, or bacteria that reach a large number of overlapping replication rounds. We found evidence that supports our hypothesis: gene order conservation of the most sensitive genes to the global program correlates significantly with the potential relevance of multifork replication in over 100 species. In addition, a recent study found two fundamental bacterial reproduction strategies, the first relying on (metabolically) efficient but slow growth and a second that relies on inefficient but fast growth (33). Of the two strategies, the latter perhaps exploits the coordination of these mechanisms. Also, correlations involving maximal growth rate should be taken cautiously as known doubling times are biased by lab controlled environments (34).

Recent studies show the important link between gene expression and gene location on the chromosome (35, 36). Indeed, the increase in gene dosage due to bacterial multifork replication appears as an added control mechanism of natural genetic circuits (37, 38). However, the relevance of genome organization goes beyond gene dosage fluctuations in fast growth (39, 40), and it may be influenced by chromosomal structure (39) and gene essentiality (41).

Our work builds on these studies to emphasize the genome-wide effect of the physiological state of the cell (the global program) on the control of gene expression and its coupling to genome organization. In fact, not only do we find that promoters that are most growth-rate dependent (at a single copy level) are located significantly close to *oriC* in *E. coli*, but also that this feature is conserved in species for which multifork gene dosage fluctuations are strongest. Therefore, we present the physiological control of gene expression as an additional aspect to consider if we are to elucidate the organization and evolutionary dynamics of the bacterial genome.

## Materials and Methods

### Promoter activity data and validating experiments

We obtained time series of optical density and promoter activity from public available data (18) that used a library of *E. coli* promoters expressing a fast-folding fluorescent protein and cloned in a low-copy plasmid (17). We considered only data of those experiments using minimal medium with 0.5% (w/v) glucose and supplemented amino acids and did not include experiments in which the strains did not grow (within one standard deviation of the mean growth curve) or whose promoter activity was constant and equal to zero.

We used this very library (*E. coli* K-12 MG1655 strain) in our validation experiments to test 3 representative promoters from each of the four response classes. The list of their names and their products’ are: *maoP* (macrodomain Ori protein), *slyD* (peptidyl prolyl *cis/trans*-isomerase and chaperone), *corA* (Ni^2+^/Co^2+^/Mg^2+^ transporter), *prmB* (50S ribosomal subunit protein L3 N5-glutamine methyltransferase), *rssB* (regulator of sigmaS), *amyA* (α-amylase), *yaaA* (peroxide stress resistance protein), *ghrA* (glyoxylate/hydroxypyruvate reductase A), *nudF* (ADP-sugar pyrophosphatase), *cpsG* (phosphomannomutase), *argQ* (tRNA) and *mutT* (8-oxo-dGTP diphosphatase). The reporter strains of these genes were retrieved from frozen stocks, plated in selective media, and grown overnight. Isolated colonies were grown overnight in the specific medium, then diluted 1/20 and pre-cultured for about 5h. Then, 96-well flat transparent plates containing 190μl of the specific medium were inoculated 1/20 with the pre-culture and added 50μl of mineral oil to prevent evaporation. Optical density (600 nm) and fluorescence (535 nm) were assayed in a Victor x2 (Perkin Elmer) at 10min intervals for ~8h (growth at 30°C with shaking).

Cultures were grown in M9 minimal medium with kanamycin (50μg/ml) to which either glucose, arabinose, lactose, glycerol or maltose was added to a final concentration of 0.5% (w/v). All five carbon media were also supplemented in the second set of experiments with amino acids to a final concentration of 0.2% (w/v), thus making 10 different nutrient conditions in total.

### Data processing and modeling

Growth rate time series were computed as the two-point finite differences of log(OD), μ(t)=Δlog(OD)/Δt, and promoter activities were computed as the two-point finite difference in time of fluorescence per OD unit, PA_*pl*_(t)=ΔGFP/Δt/OD. Balanced-growth data was computed from the mean time-series measurements of three technical replicates as the average value in a ~2h time-window during observable exponential growth. Promoter activity is in units of GFP/OD/h.

Automatic clustering was performed with the normalized growth rates and promoter activities (PA_*chr*_ and pa) with respect to their maximum. The euclidean pairwise distance was used to compute the linkage matrix with the unweighted average distance, which was then automatically divided into 50 clusters of which were rejected those with less than 2% of the sampled promoters. Clusters were then grouped together by visual interpretation resulting in the four classes presented in the main text. The robustness of the classification was tested against additional relative random noise of 5% and 10% normally distributed. The recovering rate is the mean number of genes classified as in the original classes expressed in percentage from 10 realizations. Promoter activity profiles of cluster 1 (either from pa and PA_*chr*_) were then fit to Eq.1 by means of non-linear least squares method. We fine-tuned the automatic classification and depending on the value estimated for *K_m_* we distinguish between linear (*K_m_*>3 dbl/h), saturable (0.1<*K_m_* <3 dbl/h) and constant (*K_m_*<0.1 dbl/h) profiles. The slope of linear profiles was obtained from linear least squares fits. Figure 3C-D shows the running averages of the sensitivities in a window of 10 genes.

### Definition of constitutive genes

We selected constitutive promoters as those lacking any interaction with DNA-binding transcriptional factors, even with weakly specific factors as IHF and H-NS. For this we used the regulatory network of *E. coli* downloaded from RegulonDB (42), arguably the best characterized regulatory network to date of any living organism. This gene list overlaps considerably with the set of the constitutive promoters independently identified by Genomic SELEX screening (43). Moreover, note that the promoter-specific values *V_m_* and *K_m_* can be modulated by factors that bind to RNA polymerase like (p)ppGpp. That the alarmone (p)ppGpp affects the expression of >30% of the genome of *E. coli* (5) denotes its relevance in gene expression control. However, (p)ppGpp’s role is beyond specific regulation. In fact, recent results show that it is a leading component for the proper, coordinated regulation of bacterial physiological state (5, 44), which is precisely the global program. For this reason, the expression control effects of (p)ppGpp with or without *dksA* are not considered to be part of a specific regulation, as neither does RegulonDB. In addition, that we find 3 ribosomal genes (*rpsT, rpsB* and *rpmE*) in our list highlights the fact that we focus only on transcriptional and not post-transcriptional regulation, because of the limitation imposed by the use of fluorescent reporter measurements.

### Environmental variability, R and phylogenetic distances

The classification of over 100 species depending on the variability of their environment used in this work was previously published (26). Classes of increasing environmental variability are obligate, specialized, aquatic, facultative, multiple and terrestrial. *E. coli* is found in the facultative class. From the original list, species with more than one chromosome, species that could not be found in the phylogenetic tree (see below), and species without a published unified genome in NCBI database were not considered in this study (Data set S2 in supplemental material). The importance of multifork dosage increase due to multifork replication in a given species, termed R, is obtained as the ratio of chromosomal replication time by the minimal doubling time for each bacterium. In fact, it is proportional to the maximum number of overlapping replication rounds. Values of R and the minimal doubling time were retrieved for 60 of the 100 species (27). Phylogenetic distances from *E. coli* were computed from the phylogenetic tree of (45) (Fig. S7 in supplemental material). These variables correlate with each other and specially with phylogenetic distance to *E. coli* (Fig. S8 in supplemental material). For this reason, we used partial linear correlations (27), which considers the correlation between two variables (position conservation and μ_max_ or R) controlling for a third confounding variable: phylogenetic distance in this case. The corrected values of position conservation, env, μ_max_ and R are the residuals of their respective rank correlations with phylogenetic distance. Hence, correlation between corrected values are not affected by phylogenetic inertia (Fig. 4C).

### Origins of replication and homology search

For the location of the origins of replication of most genomes, we used DoriC v7.0 (46), a database of bacterial and archaeal genomes available at http://tubic.tju.edu.cn/doric/ -- the update of September 15th, 2017). For genomes for which the origin of replication was not directly available, *Blochmannia Floridanus* and *Methanosarcina Acetivorans*, we used the web-tool that DoriC offers for its identification. The best results had expected values E=0 and E=3e-9, respectively (Data set S2 in supplemental material). Moreover, the replication terminus *ter* was set half the genome length away from the origin of replication as is done in related works. We obtained the homolog sequences (and their location) of the 708 constitutive genes of *E. coli* from Blastp, results with expected values above E=1e-3 were discarded (47). We quantified the position conservation of the half most sensitive genes to growth rate, located at *m*<0.2, for a given species as the probability of finding a smaller mean displacement in 10^4^ random selections among all homologs found at *m*<0.2 (independently of their sensitivity to growth rate). This protocol controls for a possible general conservation of genes near *oriC*, and for different numbers of homologs found in the set of species.

## Acknowledgements

This work was supported by PhD fellowship BES-2016-079127 (P.Y.) and grant FIS2016-78781-R (J.F.P.) from the Spanish Ministerio de Economía y Competitividad and the European Social Fund.

## Author contributions

P.Y. and J.F.P. designed the research, P.Y. performed the experiments, P.Y. and J.F.P. analyzed the results and wrote the manuscript.

## Supplementary Materials

Fig. S1. Comparison between two different approaches to compute promoter activity profiles.

Fig. S2. Comparison of the results of the automatic clustering algorithm when considering pa(μ) and PA_*chr*_(μ) profiles.

Fig. S3. The experimental profiles obtained from balanced growth in ten different media and time series in a single growing medium correlate only for genes in class 1.

Fig. S4. Correlation between the parameters obtained from fitting pa(μ) and PA_*chr*_(μ) to Eq.(1).

Fig. S5. Sigma factors, AT content and macrodomains do not explain the different classes.

Fig. S6. Two possible scenarios for the effect of multifork gene dosage as part of the global program.

Fig. S7. Phylogenetic tree of the 100 species used in this work.

Fig. S8. Spearman rank correlations between possible explanatory variables for conservation measures.

Data set S1. List of genes with their corresponding classes and best-fit parameters.

Data set S2. List of species with their corresponding parameters.

## Supplementary Materials

**Fig. S1.**
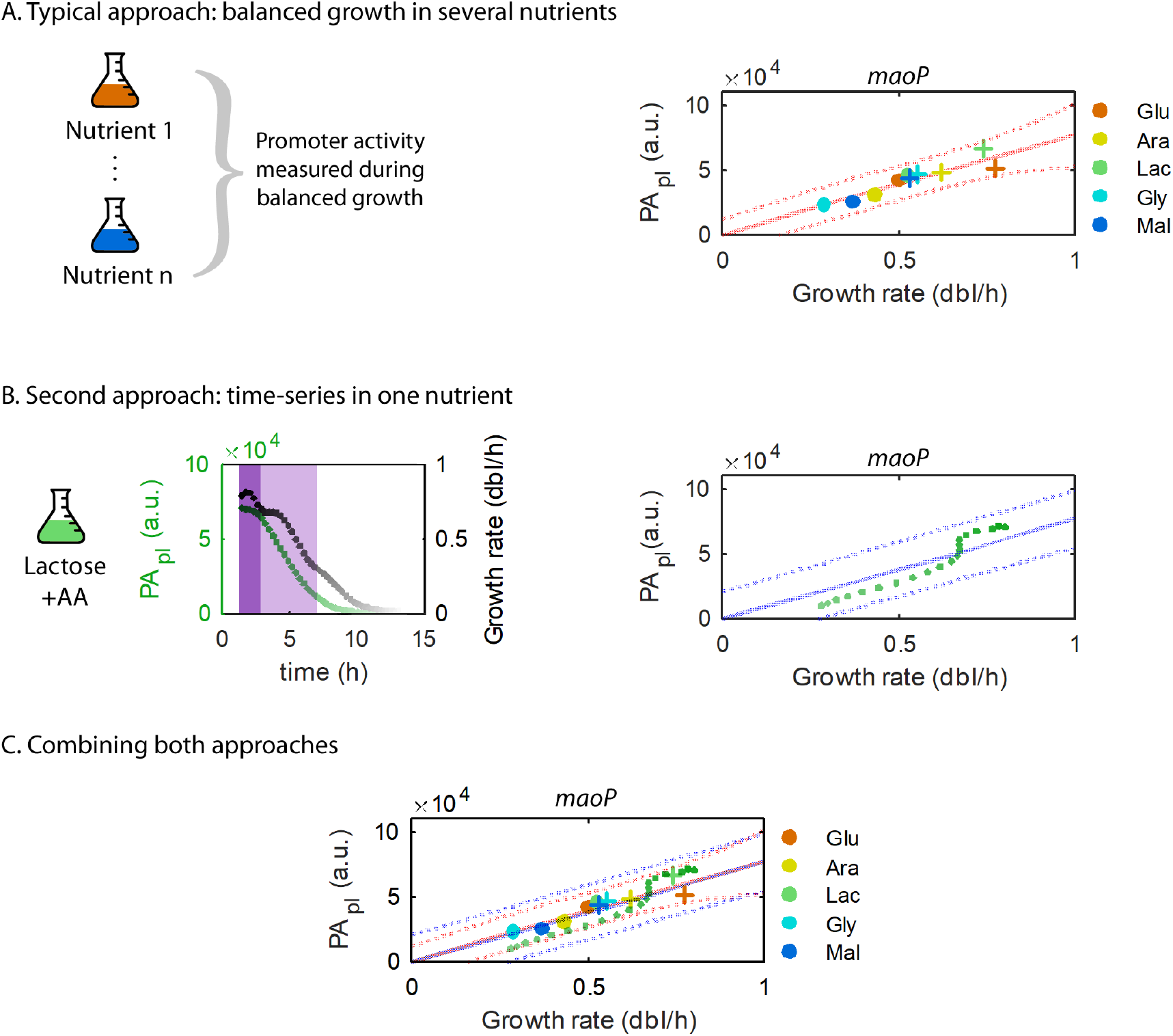
Comparison between two different approaches to compute promoter activity profiles. (**A**) The traditional approach consists in obtaining data pairs (PA_*pl*_, μ) from cultures during balanced growth in a variety of growth media. Experimental profile obtained for the promoter of gene *maoP* in minimal medium with five different carbon sources: glucose (orange), arabinose (yellow), lactose (green), glycerol (light blue), maltose (dark blue) and supplemented, or not, with amino acids (AA; crosses and circles respectively; Materials and methods). The best fit to Eq.(1) is also shown with the 95% confidence interval (red solid and dotted lines respectively). (**B**) We can also obtain promoter activity profiles from growth time-series in a single growth medium (e.g. lactose+AA). We consider promoter activity (in green) and growth rate (in black) points during early and late exponential phase (dark and light purple shade) to obtain a similar promoter activity profile (Materials and methods). The best fit to Eq. (1) and the 95% confidence bounds are also shown (blue solid and dotted lines respectively). (**C**) Superposition of the two profiles obtained previously by different means. These two approaches yield similar qualitative and quantitative profiles in this and other cases for promoters of genes in class 1.

**Fig. S2.**
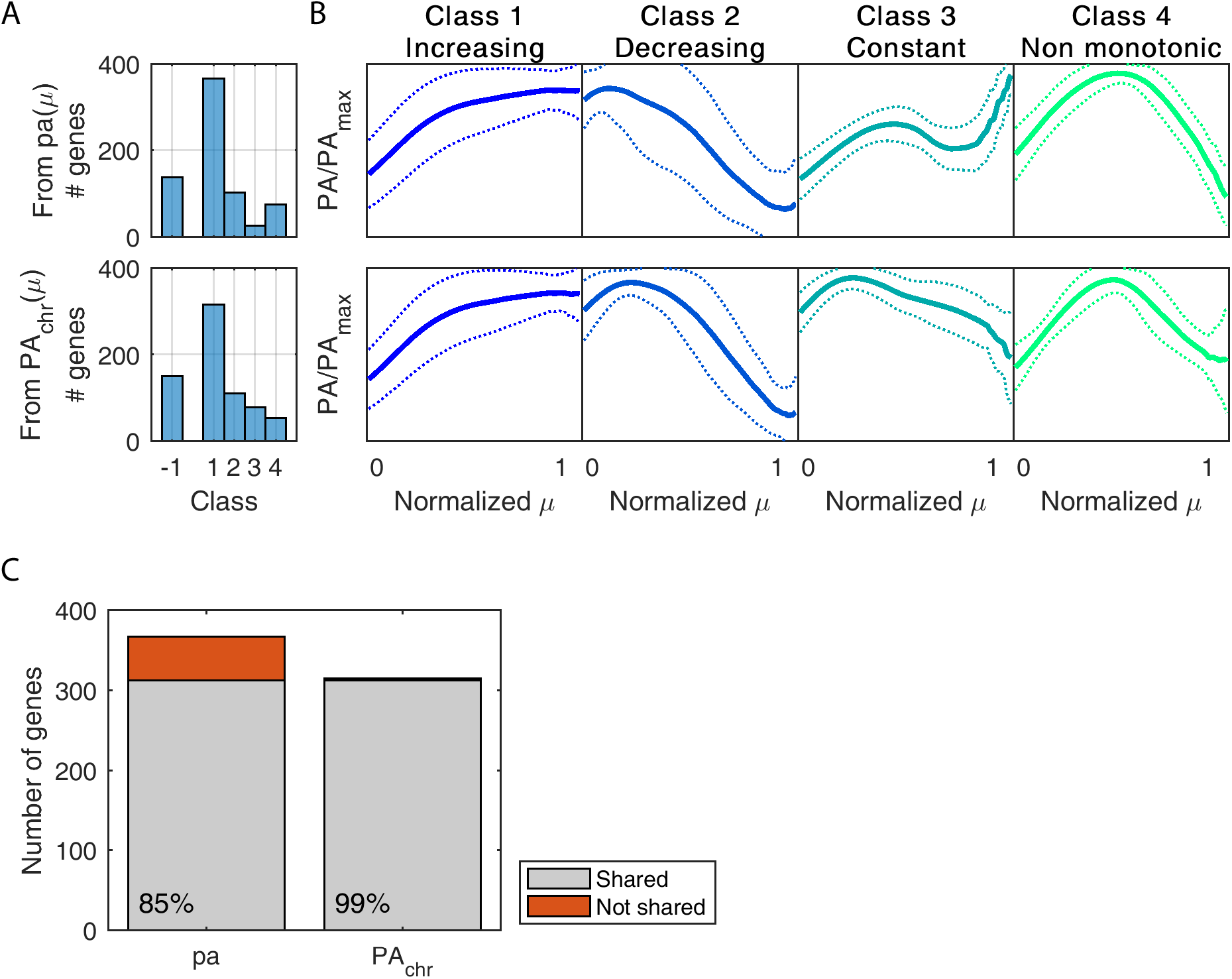
Comparison of the results of the automatic clustering algorithm when considering pa(μ) and PA_*chr*_(μ) profiles. (**A**) Histograms with the number of promoters found in each class. Class −1 includes all discarded promoters during the clustering algorithm (Materials and methods). (**B**) Mean profiles (solid lines) with one standard deviation (dotted lines) of each class. (**C**) The composition of class 1 is robust whether computed from pa(μ) or PA_*chr*_(μ) profiles. In fact, 85% (312 out of 367) and 99% (312 out of 315) of genes are shared, respectively.

**Fig. S3.**
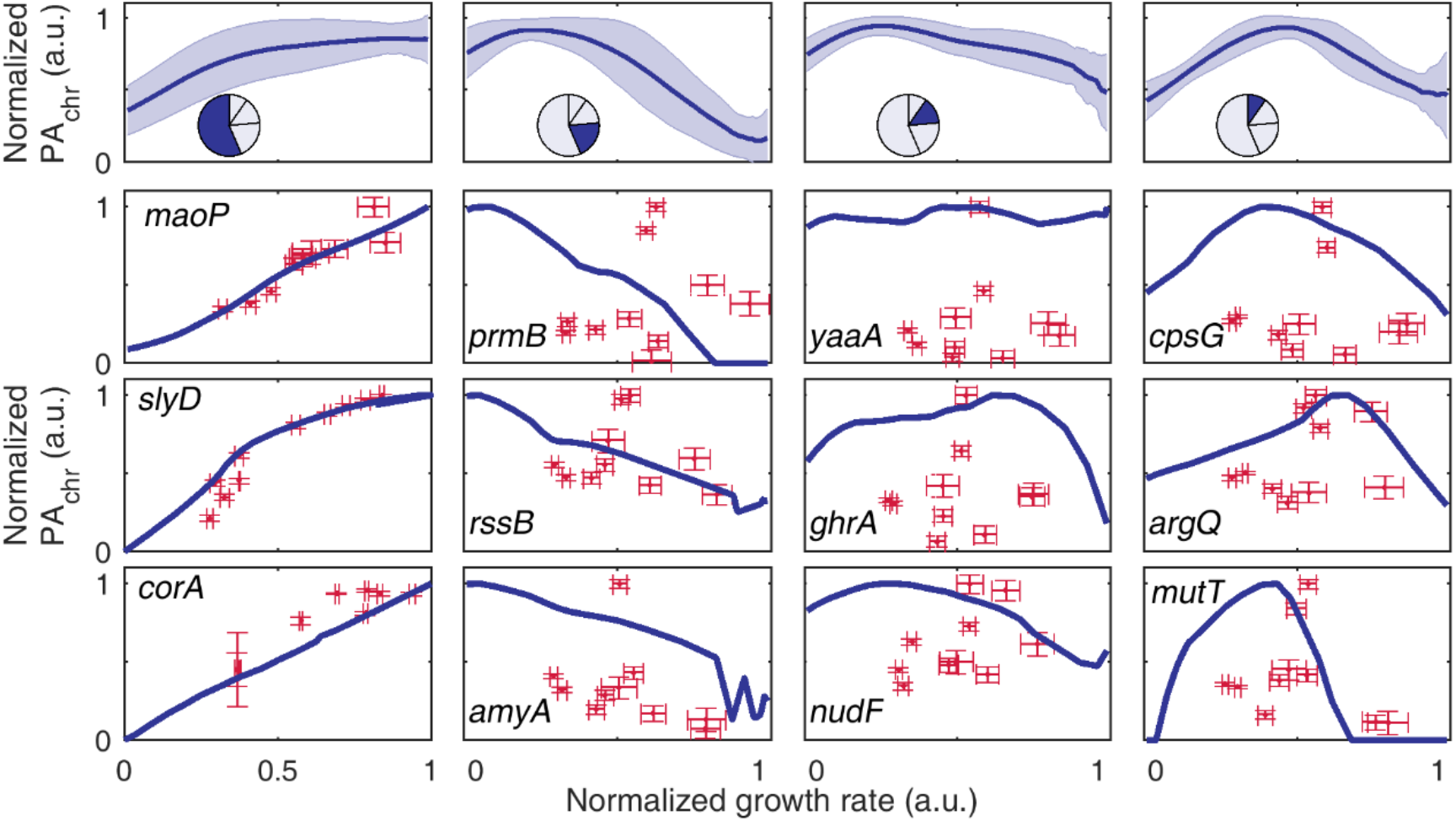
The experimental profiles obtained from balanced growth in ten different media and time series in a single growing medium correlate only for genes in class 1. (**A**) Mean PA_*chr*_(μ) profile (blue line, shaded area represents one standard deviation) of each of the four classes obtained from the clustering algorithm (Materials and Methods). Inset: pie chart with the fraction of promoters found in that particular class. (B) Experimental measurements of PA_*chr*_ from balanced growth in ten different media (red crosses, mean and sd from three replicates) of 12 genes (three from each class). Profiles from time series data in glucose supplemented with amino acids are superimposed in blue solid line [Zaslaver *et al*. (2009)]. Observe that the expected hyperbolic pattern for constitutive genes is only recovered for those in class 1. Data of *corA* grown in glycerol and arabinose resulted in fluorescence levels below the background and are not shown.

**Fig. S4.**
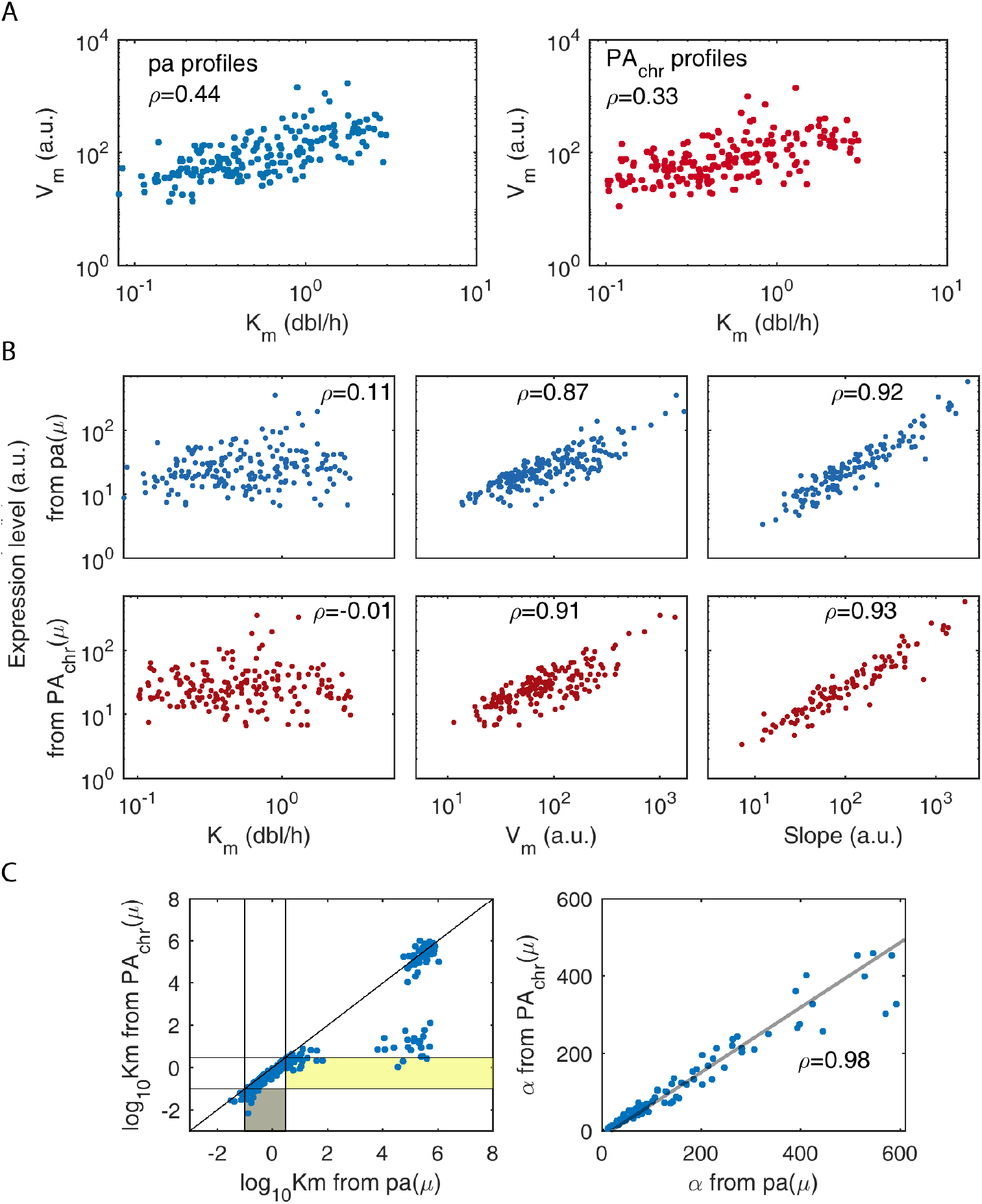
Correlation between the parameters obtained from fitting pa(μ) and PA_*chr*_(μ) to Eq.(1). (**A**) There is a small correlation between the strength of a saturable promoter (*V_m_*) and its sensitivity to growth rate (*K_m_*). (**B**) The expression level of a promoter, measured as its activity at μ=0.5 dbl/h, does not correlate with the sensitivity of saturable promoters *K_m_*. Predictably, expression level correlates well with the maximum activity of saturable promoters, and with the slope of lineal promoters. (**C**) Different sensitivities are obtained from pa(μ) and PA_*chr*_(μ) profiles. Left: values of *K_m_* of the 312 promoters belonging to class 1 when computed from pa(μ) and PA_*chr*_(μ). The region where promoters whose pa is saturable (non-saturable) but whose PA_*chr*_(μ) is constant (saturable) is marked in grey (yellow). Right: slopes of linear profiles computed from pa and PA_*chr*_ correlate linearly. In all panels, ρ is Pearson’s linear correlation coefficient, and a log-log scale is used for clarity because the values span several orders of magnitude.

**Fig. S5.**
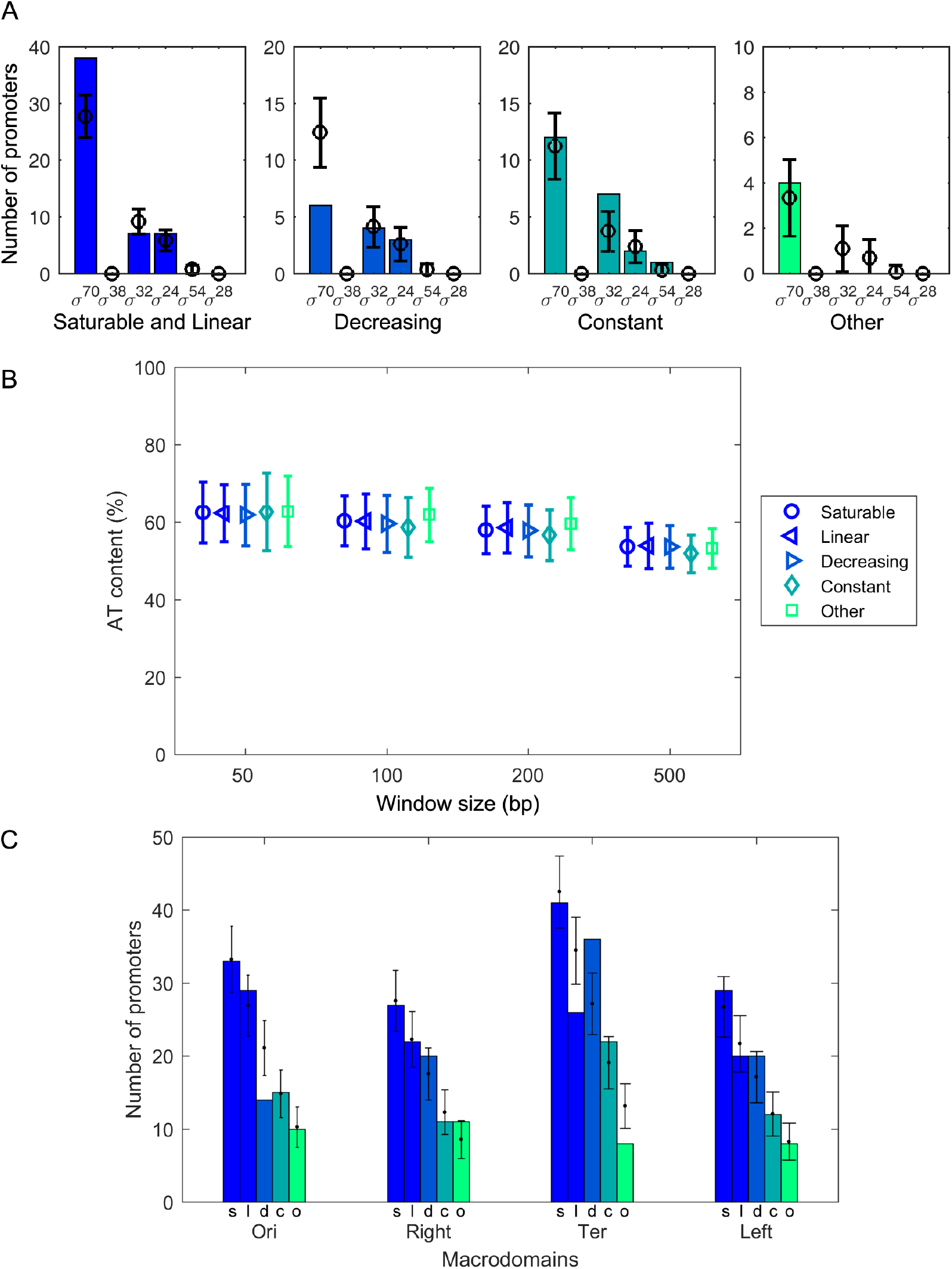
Sigma factors, AT content and macrodomains do not explain the different classes. (**A**) Number of observed genes (vertical bars) within each sigmulon (x-axis) and the expected values under 10^4^ randomizations (black circles, error bars correspond to one standard deviation; housekeeping -σ70-, general stress - σ38-, cytoplasmic stress - σ32-, extracytoplasmic stress - σ24-, nitrogen stress - σ54- and flagellar genes - σ28-). (**B**) Mean AT content and one standard deviation (y-axis) in different windows of the upstream region of the initiation transcription site (x-axis). Although constitutive promoters are often considered to be about 100bp long, we show the results of different window sizes. (**C**) Number of promoters of each class found in the chromosomal macrodomains (colored bars; classes: saturable -s-, linear -l-, decreasing -d-, constant -c-, and other -o-). Black points and error bars correspond to the mean and one standard deviation of the expected number of promoters found under 10^5^ randomizations of their location.

**Fig. S6.**
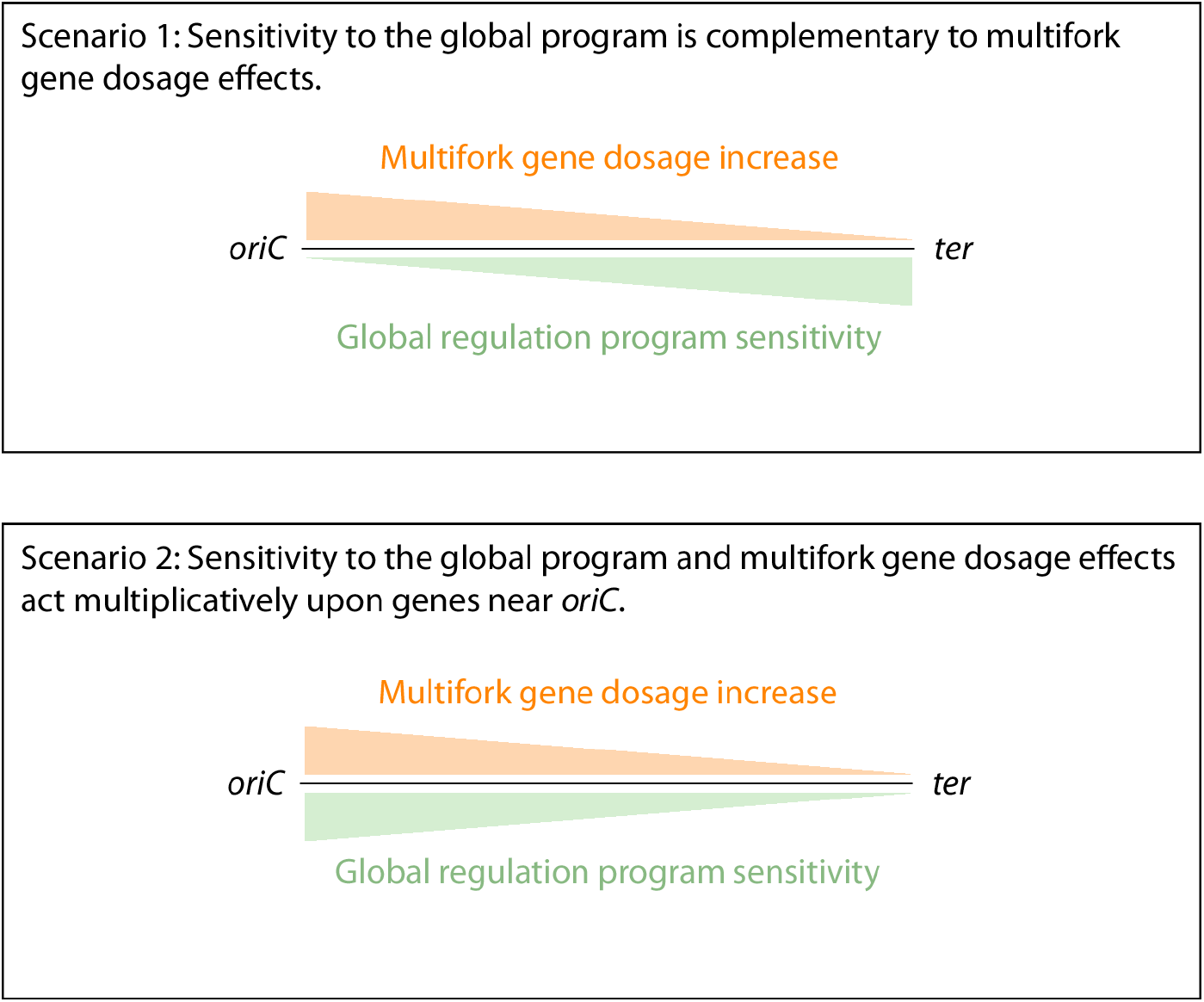
Two possible scenarios for the effect of multifork gene dosage as part of the global program. (**Top**) Scenario 1. The most sensitive genes to the global program locate near the *ter* region (symbolized by a green gradient; note that global regulation here excluded the impact of gene dosage). In contrast, the multifork effect is stronger near *oriC* (orange gradient). The gene dosage effect is consequently not coupled to a strong sensitivity. (**Bottom**) Scenario 2. The most sensitive genes to the global regulation (excluding gene dosage) locate in the *oriC* region. In this case, the effect of gene dosage is linked to the strong sensitivity.

**Fig. S7.**
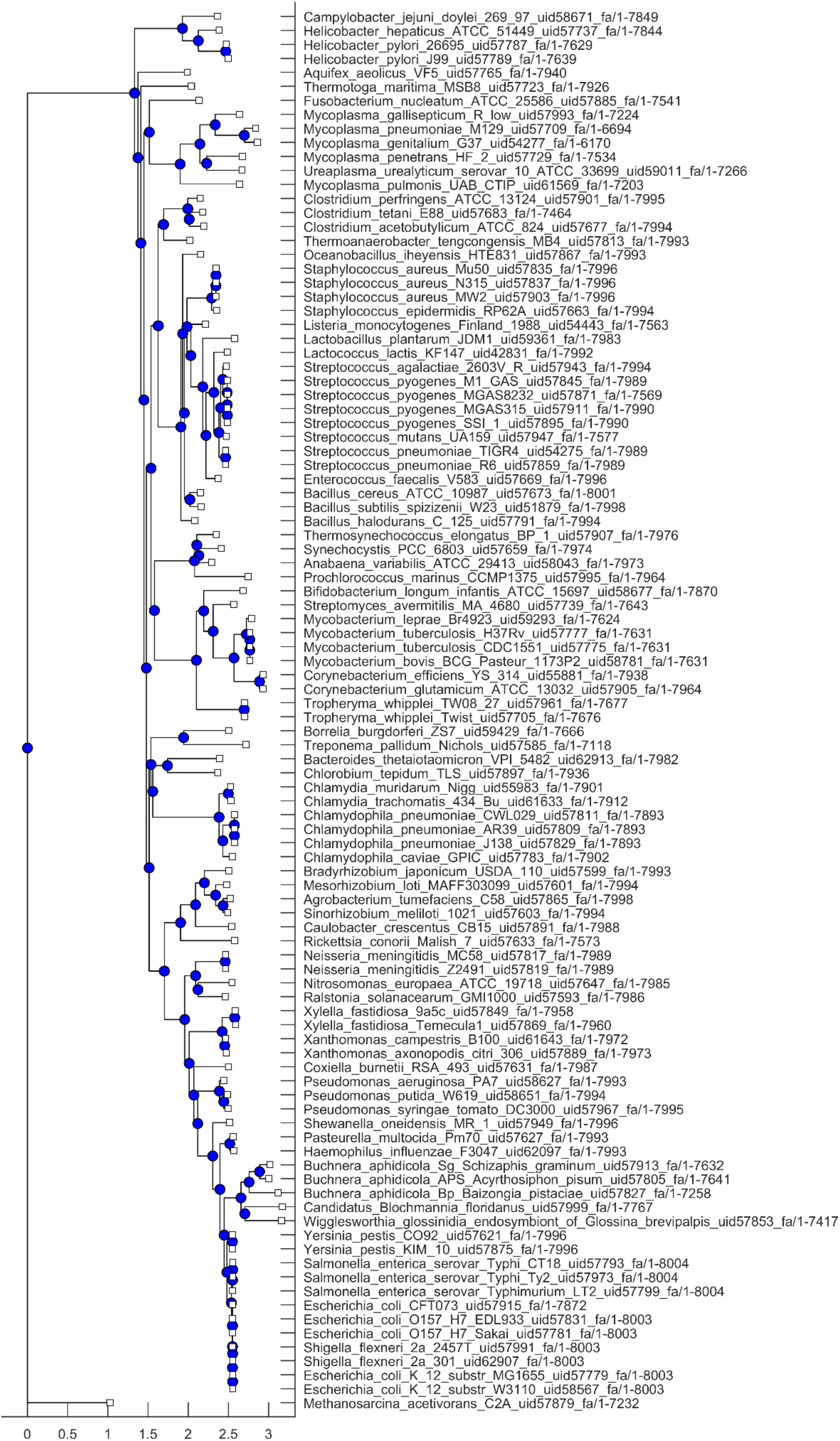
Phylogenetic tree of the 100 species used in this work.

**Fig. S8.**
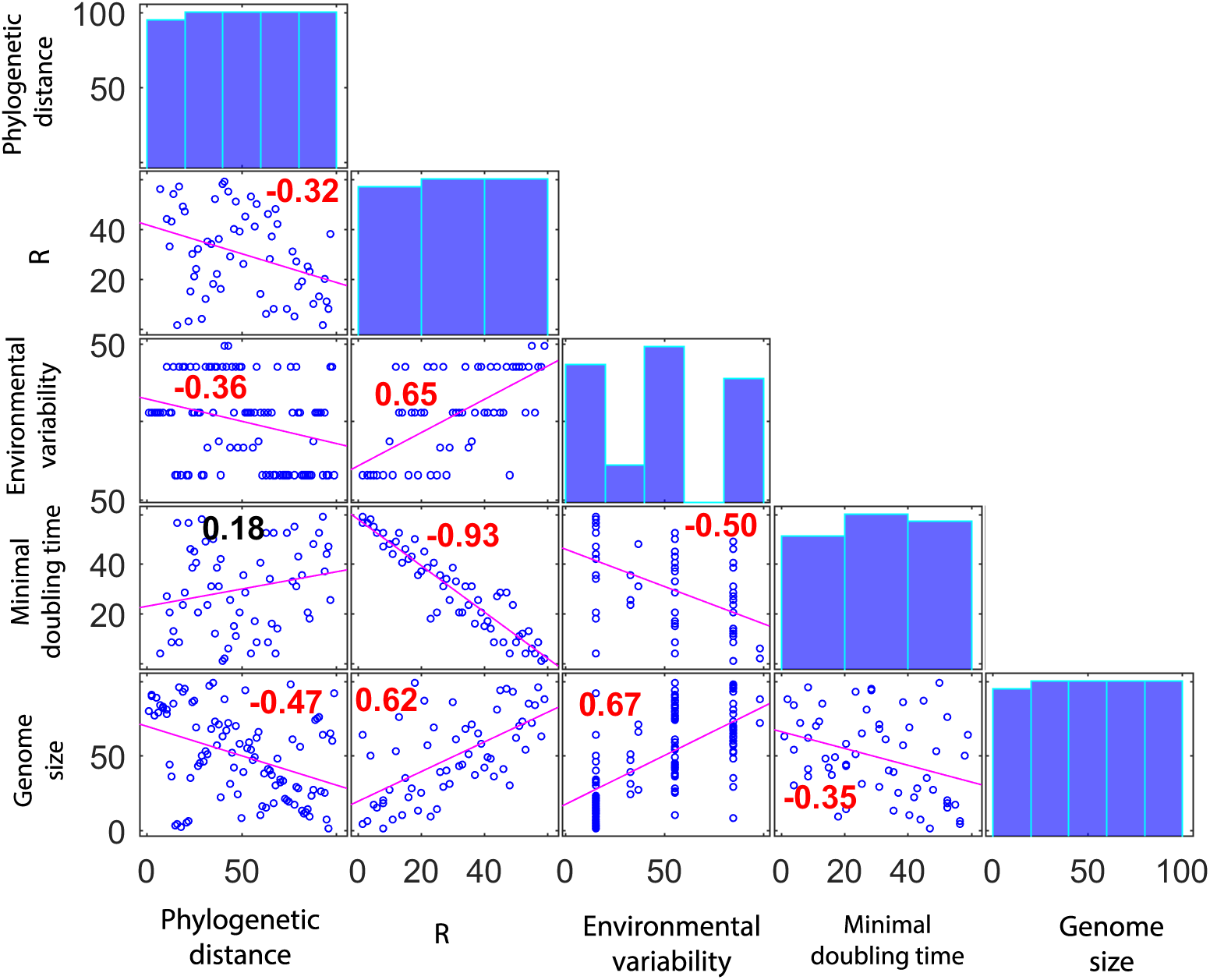
Spearman rank correlations between possible explanatory variables for conservation measures. Correlations (with ties) for all species between phylogenetic distance, R (i.e. the relevance of multifork effects), environmental variability, minimal doubling time and genome size in Mbp, when available (Materials and methods). Values in red denote significant correlations (p<0.05). Histograms of the ranks are shown in the diagonal.

## References

1. Kjeldgaard NO, Maaloe O, Schaechter M. 1958. The transition between different physiological states during balanced growth of Salmonella typhimurium. J Gen Microbiol 19:607–616.

2. Schaechter M, Maaloe O, Kjeldgaard NO. 1958. Dependency on medium and temperature of cell size and chemical composition during balanced grown of Salmonella typhimurium. J Gen Microbiol 19:592–606.

3. Kubitschek HE. 1974. Constancy of the ratio of DNA to cell volume in steady-state cultures of Escherichia coli B-r. Biophysical Journal 14:119–123.

4. Liang S-T, Bipatnath M, Xu Y-C, Chen S-L, Dennis P, Ehrenberg M, Bremer H. 1999. Activities of constitutive promoters in Escherichia coli1. Journal of Molecular Biology 292:19–37.

5. Traxler MF, Summers SM, Nguyen H-T, Zacharia VM, Hightower GA, Smith JT, Conway T. 2008. The global, ppGpp-mediated stringent response to amino acid starvation in Escherichia coli. Molecular Microbiology 68:1128–1148.

6. Klumpp S, Zhang Z, Hwa T. 2009. Growth Rate-Dependent Global Effects on Gene Expression in Bacteria. Cell 139:1366–1375.

7. Deris JB, Kim M, Zhang Z, Okano H, Hermsen R, Groisman A, Hwa T. 2013. The Innate Growth Bistability and Fitness Landscapes of Antibiotic-Resistant Bacteria. Science 342:1237435.

8. Scott M, Gunderson CW, Mateescu EM, Zhang Z, Hwa T. 2010. Interdependence of Cell Growth and Gene Expression: Origins and Consequences. Science 330:1099–1102.

9. Klumpp S, Hwa T. 2008. Growth-rate-dependent partitioning of RNA polymerases in bacteria. PNAS 105:20245–20250.

10. Peebo K, Valgepea K, Maser A, Nahku R, Adamberg K, Vilu R. 2015. Proteome reallocation in Escherichia coli with increasing specific growth rate. Mol BioSyst 11:1184–1193.

11. Weiße AY, Oyarzún DA, Danos V, Swain PS. 2015. Mechanistic links between cellular trade-offs, gene expression, and growth. Proceedings of the National Academy of Sciences 112:E1038–E1047.

12. Berthoumieux S, de Jong H, Baptist G, Pinel C, Ranquet C, Ropers D, Geiselmann J. 2013. Shared control of gene expression in bacteria by transcription factors and global physiology of the cell. Mol Syst Biol 9.

13. Gerosa L, Kochanowski K, Heinemann M, Sauer U. 2013. Dissecting specific and global transcriptional regulation of bacterial gene expression. Mol Syst Biol 9:658.

14. Kochanowski K, Gerosa L, Brunner SF, Christodoulou D, Nikolaev YV, Sauer U. 2017. Few regulatory metabolites coordinate expression of central metabolic genes in Escherichia coli. Molecular Systems Biology 13:903.

15. Camas FM, Poyatos JF. 2008. What Determines the Assembly of Transcriptional Network Motifs in Escherichia coli? PLoS ONE 3:e3657.

16. Gyorfy Z, Draskovits G, Vernyik V, Blattner FF, Gaal T, Posfai G. 2015. Engineered ribosomal RNA operon copy-number variants of E. coli reveal the evolutionary trade-offs shaping rRNA operon number. Nucleic Acids Research 43:1783–1794.

17. Zaslaver A, Bren A, Ronen M, Itzkovitz S, Kikoin I, Shavit S, Liebermeister W, Surette MG, Alon U. 2006. A comprehensive library of fluorescent transcriptional reporters for *Escherichia coli*. Nature Methods 3:623–628.

18. Zaslaver A, Kaplan S, Bren A, Jinich A, Mayo A, Dekel E, Alon U, Itzkovitz S. 2009. Invariant Distribution of Promoter Activities in Escherichia coli. PLOS Computational Biology 5:e1000545.

19. del Solar G, Giraldo R, Ruiz-Echevarría MJ, Espinosa M, Díaz-Orejas R. 1998. Replication and Control of Circular Bacterial Plasmids. Microbiol Mol Biol Rev 62:434–464.

20. Morrison PF, Chattoraj DK. 2004. Replication of a unit-copy plasmid F in the bacterial cell cycle: a replication rate function analysis. Plasmid 52:13–30.

21. Cooper S, Helmstetter CE. 1968. Chromosome replication and the division cycle of Escherichia coli Br. Journal of Molecular Biology 31:519–540.

22. Bremer H, Dennis P. 1996. Modulation of Chemical Composition and Other Parameters of the Cell by Growth Rate. E Coli Salmonella Cell Mol Biol 2.

23. Si F, Li D, Cox SE, Sauls JT, Azizi O, Sou C, Schwartz AB, Erickstad MJ, Jun Y, Li X, Jun S. 2017. Invariance of Initiation Mass and Predictability of Cell Size in Escherichia coli. Current Biology 27:1278–1287.

24. Donachie WD, Robinson AC. 1987. Cell division: parameter values and the process. Escherichia coli and Salmonella typhimurium: cellular and molecular biology American Society for Microbiology, Washington, DC 1578–1593.

25. Nanninga N, Woldringh C. 1985. Cell growth, genome duplication, and cell division. Molecular Cytology of Escherichia coli, Academic Press, London 259–318.

26. Parter M, Kashtan N, Alon U. 2007. Environmental variability and modularity of bacterial metabolic networks. BMC Evolutionary Biology 7:169.

27. Couturier E, Rocha EPC. 2006. Replication-associated gene dosage effects shape the genomes of fast-growing bacteria but only for transcription and translation genes. Molecular Microbiology 59:1506–1518.

28. Dorman CJ, Dorman MJ. 2016. DNA supercoiling is a fundamental regulatory principle in the control of bacterial gene expression. Biophys Rev 8:89–100.

29. Mitchison G. 2005. The regional rule for bacterial base composition. Trends in Genetics 21:440–443.

30. Estrem ST, Gaal T, Ross W, Gourse RL. 1998. Identification of an UP element consensus sequence for bacterial promoters. Proc Natl Acad Sci U S A 95:9761–9766.

31. Lal A, Dhar A, Trostel A, Kouzine F, Seshasayee ASN, Adhya S. 2016. Genome scale patterns of supercoiling in a bacterial chromosome. Nature Communications 7:11055.

32. Valens M, Penaud S, Rossignol M, Cornet F, Boccard F. 2004. Macrodomain organization of the Escherichia coli chromosome. The EMBO Journal 23:4330–4341.

33. Roller BRK, Stoddard SF, Schmidt TM. 2016. Exploiting rRNA operon copy number to investigate bacterial reproductive strategies. Nature Microbiology 1:16160.

34. Gibson B, Wilson DJ, Feil E, Eyre-Walker A. 2018. The distribution of bacterial doubling times in the wild. Proc R Soc B 285:20180789.

35. Block DHS, Hussein R, Liang LW, Lim HN. 2012. Regulatory consequences of gene translocation in bacteria. Nucleic Acids Res 40:8979–8992.

36. Bryant JA, Sellars LE, Busby SJW, Lee DJ. 2014. Chromosome position effects on gene expression in Escherichia coli K-12. Nucleic Acids Res 42:11383–11392.

37. Bar-Ziv R, Voichek Y, Barkai N. 2016. Dealing with Gene-Dosage Imbalance during S Phase. Trends in Genetics 32:717–723.

38. Slager J, Veening J-W. 2016. Hard-Wired Control of Bacterial Processes by Chromosomal Gene Location. Trends Microbiol 24:788–800.

39. Sobetzko P, Travers A, Muskhelishvili G. 2012. Gene order and chromosome dynamics coordinate spatiotemporal gene expression during the bacterial growth cycle. PNAS 109:E42–E50.

40. Soler-Bistué A, Timmermans M, Mazel D. 2017. The Proximity of Ribosomal Protein Genes to oriC Enhances Vibrio cholerae Fitness in the Absence of Multifork Replication. mBio 8:e00097–17.

41. Rocha EPC, Danchin A. 2003. Essentiality, not expressiveness, drives gene-strand bias in bacteria. Nat Genet 34:377–378.

42. Gama-Castro S, Salgado H, Santos-Zavaleta A, Ledezma-Tejeida D, Muñiz-Rascado L, García-Sotelo JS, Alquicira-Hernández K, Martínez-Flores I, Pannier L, Castro-Mondragón JA, Medina-Rivera A, Solano-Lira H, Bonavides-Martínez C, Pérez-Rueda E, Alquicira-Hernández S, Porrón-Sotelo L, López-Fuentes A, Hernández-Koutoucheva A, Del Moral-Chávez V, Rinaldi F, Collado-Vides J. 2016. RegulonDB version 9.0: high-level integration of gene regulation, coexpression, motif clustering and beyond. Nucleic Acids Res 44:D133–143.

43. Shimada T, Tanaka K, Ishihama A. 2017. The whole set of the constitutive promoters recognized by four minor sigma subunits of Escherichia coli RNA polymerase. PLOS ONE 12:e0179181.

44. Dennis PP, Ehrenberg M, Bremer H. 2004. Control of rRNA Synthesis in Escherichia coli: a Systems Biology Approach. Microbiology and Molecular Biology Reviews 68:639–668.

45. Lang JM, Darling AE, Eisen JA. 2013. Phylogeny of Bacterial and Archaeal Genomes Using Conserved Genes: Supertrees and Supermatrices. PLOS ONE 8:e62510.

46. Gao F, Luo H, Zhang C-T. 2013. DoriC 5.0: an updated database of oriC regions in both bacterial and archaeal genomes. Nucleic Acids Res 41:D90–D93.

47. Pearson WR. 2013. An Introduction to Sequence Similarity (“Homology”) Searching. Curr Protoc Bioinformatics 0 3.

